# Evolutionary Dynamics of microRNA target sites across vertebrate evolution

**DOI:** 10.1101/693069

**Authors:** Alfred Simkin, Rene Geissler, Alexa B. R. McIntyre, Andrew Grimson

**Author notes:** Corresponding Authors: Andrew Grimson, 445 Biotechnology Building, Cornell University, Ithaca, New York 14853., and Alfred Simkin, 124G McMichael Science Building, Campus Box 2625, Elon University, Elon, North Carolina 27244. **Author Contributions** A.S. and A.G. conceived the study. Analyses were performed primarily by A.S., with assistance from A.McI. and R.G.; A.S. and A.G. interpreted the results, and wrote the manuscript, with assistance from R.G.

## Abstract

MicroRNAs (miRNAs) control the abundance of the majority of the vertebrate transcriptome. The recognition sequences, or target sites, for bilaterian miRNAs are found predominantly in the 3′ untranslated regions (3′UTRs) of mRNAs, and are amongst the most highly conserved motifs within 3′UTRs. However, little is known regarding the evolutionary pressures that lead to loss and gain of such target sites. Here, we quantify the selective pressures that act upon miRNA target sites. Notably, selective pressure extends beyond deeply conserved binding sites to those that have undergone recent substitutions. Our approach reveals that even amongst ancient animal miRNAs, which exert the strongest selective pressures on 3′UTR sequences, there are striking differences in patterns of target site evolution between miRNAs. Considering only ancient animal miRNAs, we find three distinct miRNA groups, each exhibiting characteristic rates of target site gain and loss during mammalian evolution. The first group both loses and gains sites rarely. The second group shows selection only against site loss, with site gains occurring at a neutral rate, whereas the third loses and gains sites at neutral or above expected rates. Furthermore, mutations that alter strength of existing target sites are disfavored. Applying our approach to individual transcripts reveals variation in the distribution of selective pressure across the transcriptome and between miRNAs, ranging from strong selection acting on a small subset of targets of some miRNAs, to weak selection on many targets for other miRNAs. miR-20 and miR-30, and many other miRNAs, exhibit broad, deeply conserved targeting, while several other comparably ancient miRNAs show a lack of selective constraint, and a small number, including mir-146, exhibit striking evidence of rapidly evolving target sites. Our approach adds valuable perspective on the evolution of miRNAs and their targets, and can also be applied to characterize other 3′UTR regulatory motifs.

**AUTHOR SUMMARY:** Gene regulation typically involves two components; the first is a regulator, and the second is a target that the regulator acts on. Examining the evolution of both components allows us to make inferences regarding how gene regulation originated and how it is changing over time. One gene regulatory system, which is widespread in mammals and other animals, relies upon regulators known as microRNAs (miRNAs), which recognize target sites within mRNAs, leading to post-transcriptional repression of the targeted mRNAs. This system has been studied extensively from the perspective of the evolution of the miRNA regulators. Moreover, a subset of the target sites themselves are known to be deeply conserved. Here, we have systematically examined the rate at which target sites for individual miRNAs are created and destroyed across mammalian evolution. We find that mutations that strengthen or weaken existing target sites are strongly disfavored. For ancient microRNAs, even recently evolved target sites are under strong selective constraint, and as miRNAs age, they tend to initially experience selection only against loss of existing target sites, and later accumulate strong selection against gain of novel target sites.

## INTRODUCTION

A fundamental property of biological systems is the regulation of gene expression. In multi-cellular organisms, diverse gene regulatory systems have evolved to generate controlled gene expression patterns across different tissues, and in response to different environments. As organismal complexity increases, the complexity of regulatory networks and mechanisms appears to have increased at a rate that is greater than any increase in the number of genes [1–3]. Thus, understanding the evolutionary pressures acting on different regulatory pathways is central to our understanding of the biology of complex organisms.

A major driver of evolutionary change is differential control of common gene complements. For example, humans and chimpanzees share 97% of their genomes [4], and yet exhibit a large number of strikingly divergent anatomical and behavioral traits. Primates, nematodes, and flies have similar numbers of protein-coding genes, many of which are conserved in their primary sequence [5], and yet the three clades exhibit marked phenotypic differences. Different species of cichlid fish that shared a common ancestor within the past one million years, and which are virtually identical genetically, display a remarkably diversified set of morphological and behavioral traits [6].

Changes, including those that appear relatively modest, in gene regulatory networks underlie many of the most striking observed phenotypic and developmental differences between life forms, and the study of these changes is essential to a complete understanding of the evolution of biological systems.

Regulatory networks are mediated by *trans* acting molecules (regulators) that interact with target genes or transcripts by way of *cis* recognition sites found on the targeted gene or gene product.

Changes in gene regulatory networks can thus occur by two main mechanisms: either the *trans* factor changes in such a way that interactions with all or many of the cognate *cis*-acting targets are modified, or individual *cis* targets change to favor greater or lesser recognition by a static *trans*-factor, such that the functional targets of this static *trans* factor gradually shift over time. While changes in *trans*-acting factors can quickly change entire gene regulatory networks, gradual shifts in *cis*-acting targets may lead to significant changes in the function of a *trans*-acting factor without altering *trans*-factor sequence or eliminating the general tendency for some *cis*-acting target sites to be deeply conserved. Studying the rate at which *cis*-acting sites are lost and created is thus of equal importance to studies of the rate at which *trans*-acting regulatory sequences change their binding affinities.

The evolution of gene regulatory systems is understood best for transcriptional regulation, which is mediated by *trans*-acting transcription factor proteins and their corresponding *cis*-acting binding sites in DNA. Transcription factor affinities for their target sequences tend to be highly conserved [7], yet there are also examples of factors whose binding affinities change relatively rapidly over time [8–10]. Some transcription factors are extremely conserved [7], while others appear to be functional despite their recent evolutionary origins [11]. A similar process can be observed for *cis*-acting target sites; the rate at which evolutionarily conserved *trans*-acting transcription factors acquire novel *cis*-acting target sites or lose existing target sites is referred to as transcription factor binding site (TFBS) turnover [12]. Numerous approaches have been developed to model TFBS turnover, and have made significant contributions to our understanding of regulatory networks [13–15]. It is notable that some evolutionarily ancient transcription factors exhibit high levels of TFBS turnover [16], indicating that these deeply conserved transcription factors gain and lose targets relatively frequently. This finding demonstrates that the evolutionary age of *trans* regulators and their respective populations of *cis* targets are not always well correlated.

The approaches used in the study of transcriptional regulation by transcription factors can be applied to other regulatory systems. MicroRNAs (miRNAs) are a class of small regulatory RNAs found in almost all animal species, and many other organisms [17, 18]. In bilaterian animals, miRNAs promote mRNA degradation by recognizing ∼6-8 basepair target sites, which are typically found within the 3′ untranslated region (3′UTR) region of mRNAs [19]. Thus, miRNAs represent a system of post-transcriptional gene regulation that largely follows the *trans*-*cis* paradigms established for transcription factors.

Following the discovery of miRNAs, the basic evolutionary characteristics of these regulatory molecules were studied extensively. Researchers discovered that certain miRNAs and a subset of their targets are conserved across deeply diverged phyla [20, 21], while others appear to be genus and even species specific [22]. Using techniques similar to those applied to the study of transcription factors, models have been developed to track evolutionary changes in miRNAs [23–26]. By documenting the origins and decay of novel miRNAs, evolutionary studies of changes in these *trans*-acting factors have revealed the birth and death of regulatory sequences that impact hundreds of target genes at once [27–29].

The acquisition and destruction of *cis*-acting target sites of statically defined miRNAs (miRNA target turnover) has also been studied to some extent. Early studies revealed that some conserved miRNAs have deeply conserved targets and that the target sites of miRNAs tend to be themselves deeply conserved [30–33]. Analyses of conserved miRNA target sites have been used as a metric of functionality to estimate that the majority of human protein coding genes harbor functional target sites for miRNAs in their 3′UTRs [34]. Population genetics models have been used to demonstrate that a fraction of miRNA target sites are likely to be under purifying selection [20], and in cichlid fish some studies have found that miRNA target sites appear to be under positive selection [35], although this conclusion has been contested [36].

While previous studies have demonstrated that miRNAs and their targets are under strong selective constraint, to date there are few studies [37–40] that have extended a study of turnover rate dynamics to miRNA target sites. Despite the demonstrated insights gained through examining turnover rates for transcription factor target sites [13–15], we know of few methods that have systematically examined turnover dynamics of the targets of individual miRNAs. Here, we have developed an approach that is capable of determining rates of miRNA target site turnover for individual miRNAs, and of quantifying the relative strength of selection acting on individual targets. We use this method both to characterize the evolutionary dynamics of the targets of specific miRNAs, and to further develop generalizable principles regarding the strength, prevalence, and nature of natural selection acting on the targets of miRNAs. Our results identify remarkably different patterns of target site gain and loss between different groups of miRNAs. We further show that many of the specific patterns of miRNA target site evolution we find in vertebrates are also observed in insect phylogenies.

## RESULTS

### Simulating microRNA target site evolution

To model evolutionary changes in miRNA target sites, we developed a simulation-based approach to measure expected rates of nucleotide substitution events in the absence of selection, with the goal of using this as a benchmark to compare against the observed evolution of miRNA target sites. We selected ten representative mammalian species, chosen to be evenly distributed across the mammalian phylogeny, with genome sequences in at least their second revision, and with human and mouse representing the most distantly related species (Fig 1A). We reconstructed ancestral sequences using DNAML [41], which we used to estimate two parameters. First, by comparing inferred ancestral 8mer sequences to descendant 8mer sequences, we were able to directly infer changes in miRNA binding sites in the presence of natural selection. Second, by recording the substitution frequencies for every nucleotide given both of its flanking nucleotides, we were able to infer nucleotide substitution probabilities, which enabled us to create simulated sequences with randomly placed substitutions that account for the impact of flanking nucleotides on sequence evolution (Fig 1B). The simulated dataset, therefore, was comprised of 3′UTRs with randomly placed substitutions, and served as a proxy for neutrally evolving 3′UTRs. We compared this simulated dataset against real 3′UTR sequences (observed), and their DNAML inferred ancestors. This approach allowed us to compare miRNA target site losses and gains in the observed dataset to those seen in sets of simulated neutrally evolving sequence.

**Fig 1.**
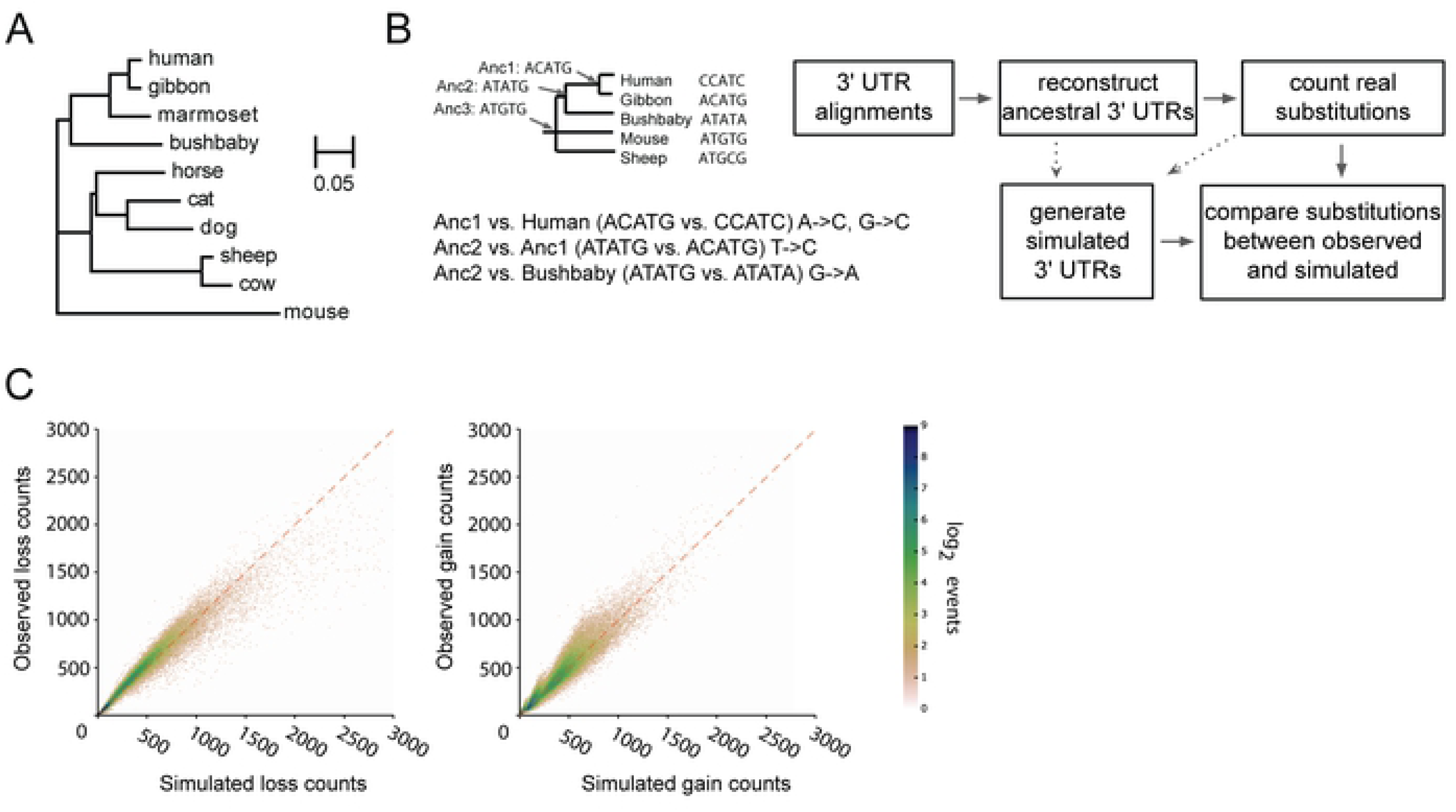
Simulated 3′UTR sequences recapitulate patterns of sequence evolution observed for sequence motifs. **A)** Mammalian phylogeny used in this study. **B)** Schematic showing how ancestral nodes of a phylogeny are generated, and example comparisons between ancestral nodes and their direct descendants (left). Flowchart of experimental pipeline (right). **C)** Comparison of loss (left) and gain (right) of each possible 8mer across 3′UTR sequences, summed across species detailed in panel A, compared between simulated (x-axis; values are mean of 100 simulations) and observed (y-axis). Count densities (see log_2_ scalebar), have a pseudocount of 1 added, and represent the number of points within a distance of 3 from the point in question. Dashed red line indicates an idealized x=y relationship between simulated and observed counts.

The primary determinant by which miRNAs recognize their target sites is through basepairing between the 5′ end of the miRNA (seed region) and a short 6-8 basepair target site [19]. Therefore, we began by examining our observed and simulated datasets with regard to 8 nucleotide motifs (8mers) to determine whether our simulations served as an effective model of neutral sequence change in 3′UTRs. For each 8mer, we counted the number of times that it was disrupted by point substitutions and the total number of times that the 8mer arose through point substitution, which we refer to as loss and gain rates, respectively. This examination was performed for each of the 65,536 possible 8mers, and repeated using 100 independent sets of simulated 3′UTRs across the ten species. When we compared the rates of loss and gain of 8mers between our set of simulated 3′UTRs and observed 3′UTRs, we found that the mean of our simulated rates was highly correlated with the observed rates (Pearson rank correlation, R^2^=0.93 and 0.91, for losses and gains, respectively, P <10^−50^; Fig 1C). Thus, with high consistency, 8mers that are predicted to undergo rapid substitution, as modeled in the simulated 3′UTRs, correspond to those undergoing the most rapid substitution rates in reality, and vice versa. We concluded that our model accurately captures substitution rates for the majority of 8mers, and that most of the fluctuation in observed 8mer substitution rates can be accurately predicted by modeling variations in the underlying substitution rates of constituent nucleotides.

### Selection against loss and gain of miRNA target sites of ancient miRNAs

Having established that most 8mers in the observed dataset are lost and gained at rates that are equivalent to their simulated counterparts, consistent with their evolution under neutrality, we next sought to examine the loss and gain rates of 8mers corresponding to miRNA target sites. We concentrated on 8mer miRNA target sites rather than smaller sites, because as a group, 8mer sites exhibit the most pronounced evolutionary signatures in animals [19], presumably reflecting their increased efficacy. We examined first the target sites of the 75 most deeply conserved miRNAs: those miRNAs whose 8mer ‘seed’ region (the region known to basepair directly with 3′UTRs) sequences are identical in human, mouse, and zebrafish miRNAs. We refer to these miRNAs in the remainder of this text as ‘deeply conserved’. For each 8mer target site, we compared the observed loss or gain count relative to the mean of the simulated values, recording the number of standard deviations that the observed values deviated from the simulated. We then recorded the ranks of these standard deviation comparisons relative to all other 8mers (S1 and S2 Tables). We found that the target sites corresponding to the majority of the deeply conserved miRNAs (Fig 2A, in red) were lost far less frequently than expected absent selection, as judged by a comparison to all other 8mers (Fig 2A, in blue; Wilcoxon Rank test; P=7.5×10^−34^). These results confirm that our technique has power to detect previously reported strong selection against loss of existing miRNA target sites [42]. It is important to note that our approach to determining selection against target site loss is distinct from previous approaches. Conventional approaches [33, 34, 43] capture selection that maintains conserved sites, examining the lack of change that occurs on the modest subset of sites that are under selective maintenance, a concept that contributes to miRNA target site prediction algorithms. Such approaches also often use several control kmers as a proxy for neutrally evolving versions of a kmer of interest. Our approach considers the observed rates at which all target sites— not just those that are conserved—are changing across an entire phylogeny, and compares these rates against explicitly modeled expected rates of change for that same kmer motif.

**Fig 2.**
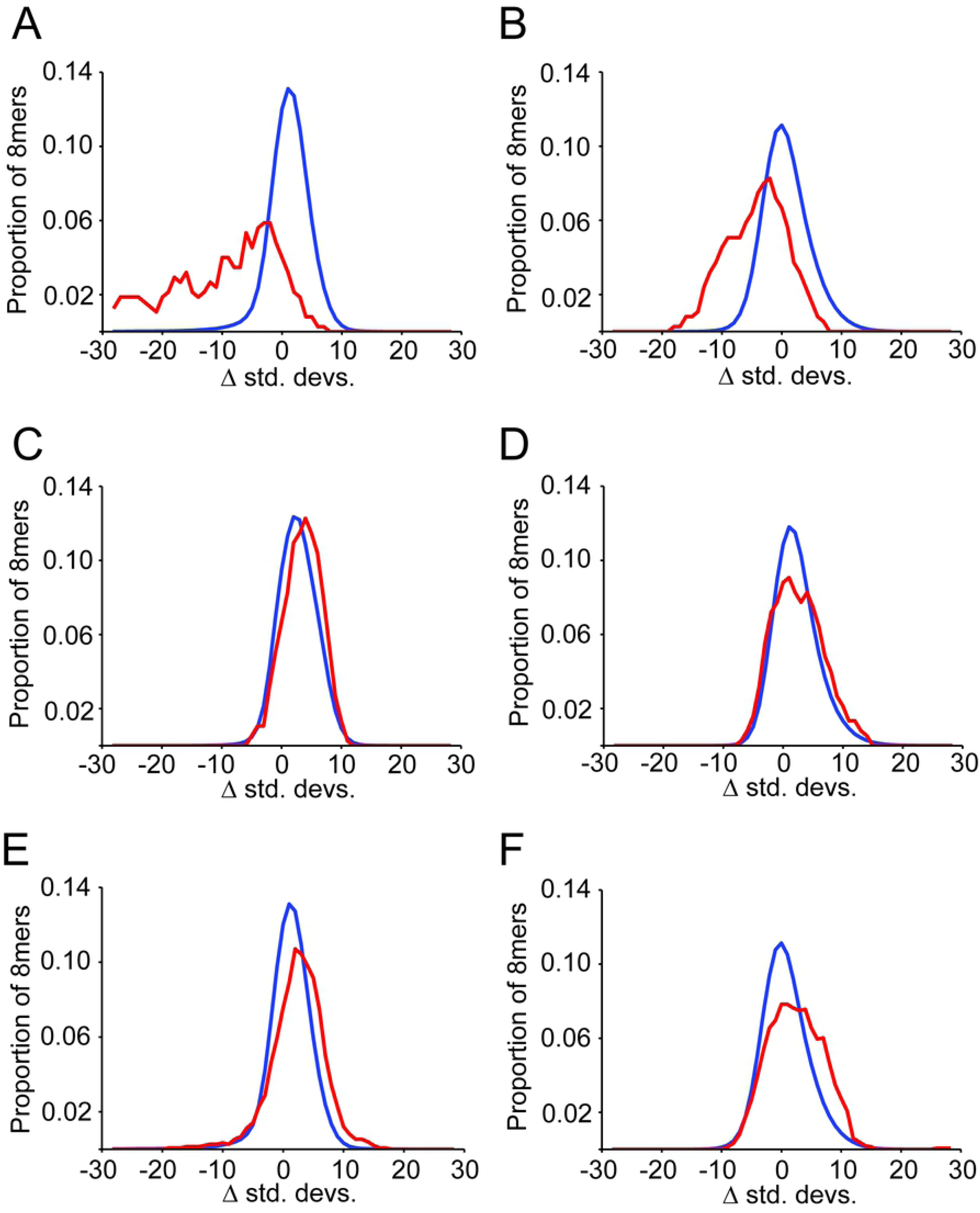
Target sites of deeply conserved miRNAs are lost and gained rarely in 3′UTRs. Each panel depicts the proportion of all 8mers (y-axis) that fall at a given number of standard deviations above or below the mean of simulations (x-axis). All 65,536 8mers are plotted in blue, and 8mer miRNA target sites in red. **(A, B)** Analysis of miRNA target site loss and gain events (respectively) in 3′UTRs, for deeply conserved miRNAs. **(C, D)** Analysis of miRNA target site loss and gain events (respectively) in intronic sequence, for deeply conserved miRNAs. **(E, F)** Analysis of miRNA target site loss and gain events (respectively) in 3′UTRs, for moderately conserved miRNAs.

In addition to characterizing selection acting against the loss of existing binding sites, our approach also allows us to determine whether novel target sites are gained at neutral rates, an important additional aspect of miRNA target site evolution. We observed strong selection against the gain of binding sites for deeply conserved miRNAs (Fig 2B, in red) relative to all other 8mers (Fig 2B, in blue; Wilcoxon Rank test; P=1.5×10^−18^; S2 Table S2). Thus, gain of target sites for deeply conserved miRNAs is sufficiently deleterious to be observable across all 3′UTRs, despite the fact that many mRNAs are not coexpressed with their cognate miRNAs, and therefore expected to evolve neutrally. It is worth noting that deviations in observed gain rates, compared to simulations, were significantly less pronounced than loss rates (P=0.012), indicating that for deeply conserved miRNAs, creating novel targets appears to be somewhat less deleterious than destroying existing targets.

To verify the specificity of our results, we examined patterns of loss and gain of miRNA target sites in a neutrally evolving region of the genome that does not participate in miRNA-mediated regulation. MiRNAs are active predominantly in 3′UTRs; we therefore chose to examine the central regions of introns as a negative control, reasoning that by omitting the canonical splice sites at intron borders, the remaining sequence is predominantly evolving at a near-neutral rate [44] and is unlikely to participate in miRNA targeting. In this intronic dataset, we observed that target sites for deeply conserved miRNAs lose and gain miRNA target sites at approximately neutral or slightly above neutral rates (Fig 2C and 2D, S3 and S4 Tables, Wilcoxon Rank tests of 0.008 and 0.518, respectively). Because losses and gains of miRNA target sites are only constrained in 3′UTRs, we conclude that our model has strong power to identify natural selection in motifs corresponding to miRNA target sites.

Returning to our 3′UTR dataset, we also examined target sites corresponding to less deeply conserved miRNAs, defined as those present both in humans and mice – the most distantly related species in our alignment – but absent from zebrafish, which we refer to as moderately conserved miRNAs. Importantly, deeply conserved and moderately conserved miRNAs are both present throughout the entire mammalian phylogeny (Fig 1A) used to examine target site evolution. When considered as a set, these moderately conserved miRNAs showed evidence of marginally faster than expected target site turnover (Fig 2E and 2F, S5 and S6 Tables, Wilcoxon Rank tests of 5.327×10^−12^ and 7.998×10^−12^, respectively). That is, rates of target site loss and gain for moderately conserved miRNAs are above those of most 8mers. It should be noted that more miRNAs were defined as moderately conserved than were defined as deeply conserved (299 versus 75) affording an increase in power to detect subtle selection acting on moderately conserved miRNAs as a group over deeply conserved miRNAs. Moderately conserved miRNAs therefore do not appear to exert detectable selective constraint against mutations within their targets when considered collectively, despite their origin prior to the common ancestor of our mammalian phylogeny.

To gain additional perspective on the extent to which deeply conserved miRNAs influence 3′UTR sequence evolution, we quantified the number of target site gains and losses observed, compared to their expected values estimated from simulated 3′UTRs. For miRNA target site losses, aggregated across the 75 deeply conserved miRNAs, we observed only 66% as many losses as expected (S1 Table). For site gains, we observed 81% as many events as expected (S2 Table). Collectively, this reduction in losses and gains corresponds to 2.0 fewer changes in miRNA target sites per 3′UTR than expected (1.5 fewer changes per site than expected for losses, and 0.5 fewer gains per site than expected, S1 and S2 Tables). Thus, for deeply conserved miRNAs, there is an extensive evolutionary signature of target site constraint during mammalian evolution, which cumulatively amounts to tens of thousands of selective events.

### Recently created miRNA binding sites are rarely mutated

It is well-established that miRNA target sites are among the most deeply conserved sequence motifs within mammalian 3′UTRs [32–34]. Yet, for deeply conserved miRNAs, their conserved target sites represent a small fraction of the sites capable of responding to their cognate miRNAs when coexpressed [30]. Importantly, the extent to which more recently evolved miRNA target sites contribute to global signatures of selection has not been thoroughly investigated. To determine whether the reduction in rates of target site gain and loss we observed (Fig 2A, 2B) is derived solely from selection acting on deeply conserved sites, we masked regions of the genome that are perfectly identical across all included species (Fig 1A) from our analysis, considering only the 8 nucleotide regions that have undergone substitutions. When we compared all 8mers containing at least one substitution against deeply conserved miRNAs whose target sites contain at least one substitution, we found that target sites of deeply conserved miRNAs were still undergoing loss and gain substitutions at rates significantly below other 8mers (Fig 3, losses and gains in parts A and B, respectively, Wilcoxon Rank test; P=4.2×10^−22^ and 2.4×10^−15^, respectively). This signature corresponds to thousands of constrained sites (S7 and S8 Tables). Importantly, these results indicate that purifying selection operates on miRNA target sites that are actively undergoing substitutions within mammals.

**Fig 3.**
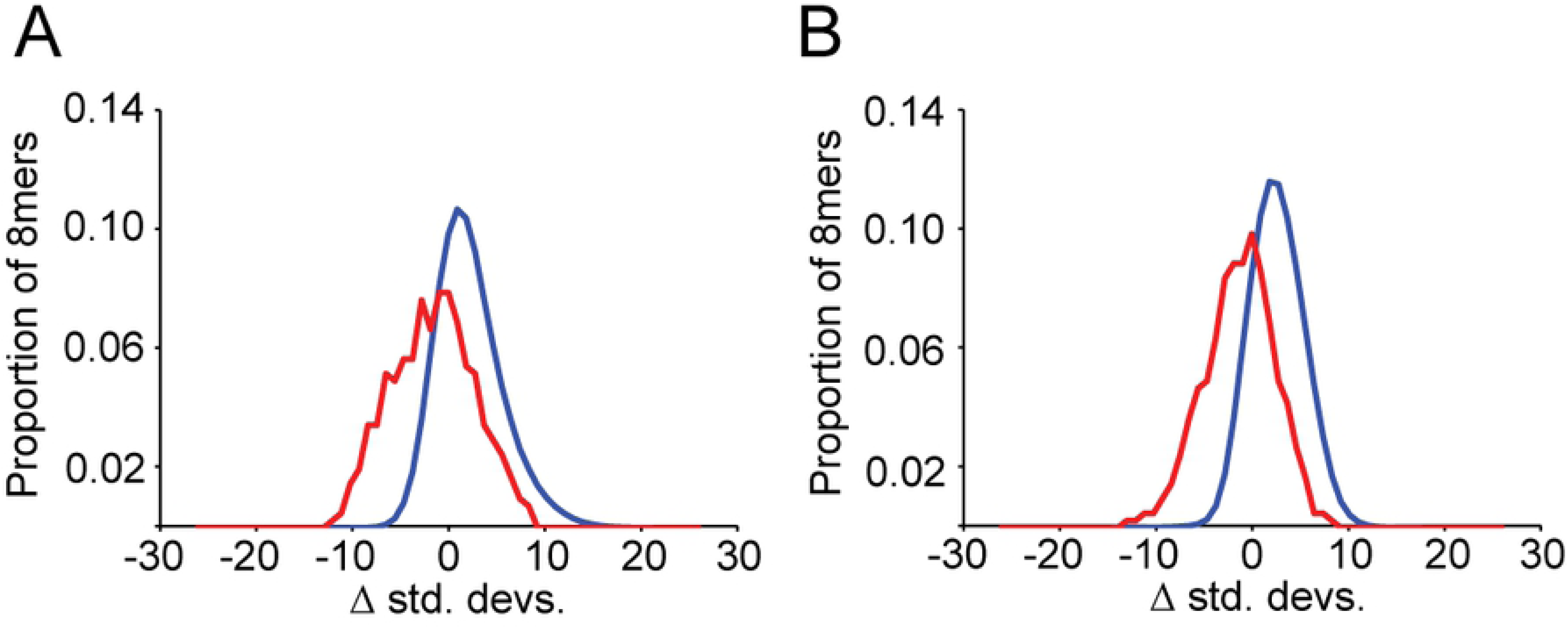
MiRNA target sites with recent substitutions are under strong selection. Analyses of the subset of target sites that have undergone at least one substitution within mammals are depicted (**A, B**; losses and gains, respectively), both for all 8mers (in blue) and for targets of deeply conserved miRNAs (in red), otherwise as described in Fig 2.

### MicroRNA age correlates with selective constraint acting on target site turnover

Having established that target sites of deeply conserved miRNAs exhibit strong selection against target site turnover, and that target sites of moderately conserved miRNAs do not exhibit strong selection, we systematically investigated how miRNA age correlates with target site turnover. We used a 100way vertebrate phylogeny [45] to determine the age of each miRNA (measured as summed branch length). We found that older miRNAs have lower loss and gain rates of target sites than younger miRNAs (Fig 4A, Fig 4B, respectively; P=9.5×10^−27^ and 6.7×10^−15^), although the degree of correlation was modest (R^2^=0.33 and 0.19, respectively). Because more ancient miRNAs tend to exhibit more constrained rates of target turnover, our observations are consistent with a model in which miRNAs begin with a relatively small subset of functional targets, and progressively acquire targets during evolution that then drive conservation of both targeting and primary miRNA sequence, consistent with results seen by others [38]. However, we also observed notable examples of miRNAs that are ancient, whose targets are gained and lost at approximately neutral rates. For example, miR-146 is deeply conserved, and yet its target sites are both lost and gained more quickly than 88% of all other 8mers. Despite being ancient, such miRNAs appear to be rapidly evolving new target genes, as has been reported previously for the rapidly evolving targets of some classes of deeply conserved transcription factors [16]. While our data provides direct evidence in support of the canonical view that conserved miRNAs generally demonstrate conserved targeting [42], our technique also identifies outlier miRNAs that are deeply conserved with rapidly evolving targets.

**Fig 4.**
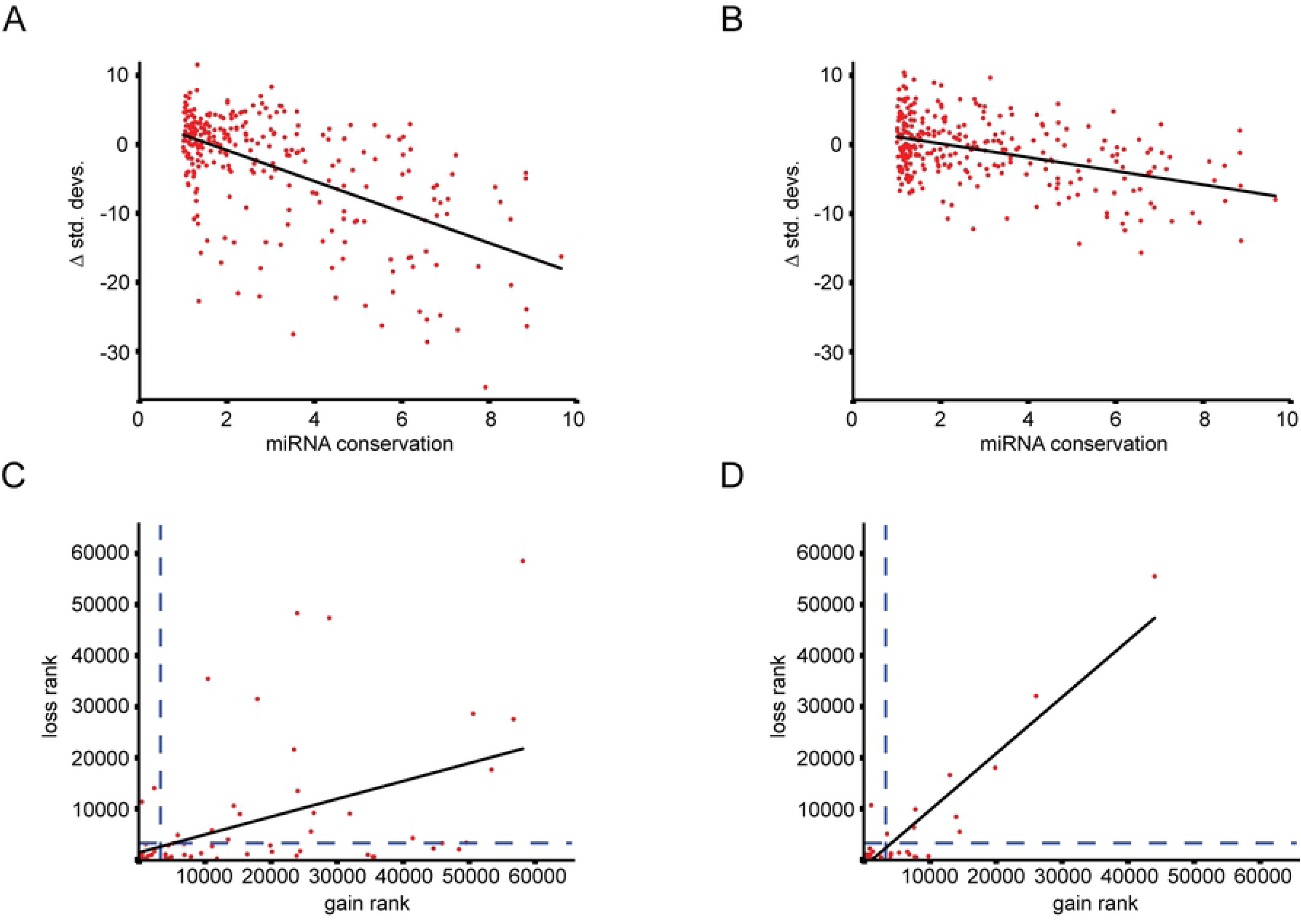
Selection against gain and loss of miRNA target sites correlates with miRNA age. The age of each miRNA (x-axis, substitutions per site), as measured by the summed branch length of all known species containing a given full length miRNA, compared against the strength of selection acting on each target site (y-axis, standard deviations below the mean of simulated values), compared for losses and gains of sites (**A and B**, respectively). **(C, D)** miRNA gain ranks relative to all 8mers (x-axis, ranked by standard deviations above or below the mean of simulations) versus miRNA loss ranks relative to all 8mers (y-axis). miRNAs considered to be under negative selection are those below a rank of 3277 (5% of 65,536, dashed lines). Relationships between loss and gain analyzed for mammalian miRNAs using 75 deeply conserved vertebrate miRNAs (C), and drosophila miRNAs using 43 deeply conserved insect miRNAs (D).

We next examined whether the miRNAs with the strongest selection against gain of deleterious target sites were also those with the strongest selection against loss of functional sites, focusing on our set of 75 deeply conserved miRNAs. We observed that many of the miRNAs with the strongest selection against gain of deleterious target sites were also those with the strongest selection against loss of existing target sites, and that there is a significant correlation between target site gain rates and target site loss rates (P=7.8×10^−6^ Fig 4C). Nonetheless, many miRNAs showed limited association between loss and gain rates, decreasing the strength of the overall correlation (R^2^=0.24). We classified each miRNA by considering a miRNA to have constrained shifts in targeting if its standard deviation score fell within the lowest 5% of all 8mers. Using this system to classify target site gain and loss rates as either ‘constrained’ or ‘unconstrained’ created four broad categories: constrained for both losses and gains, constrained for neither losses nor gains, constrained for losses but not gains, and constrained for gains but not losses. Within the 75 deeply conserved miRNAs, the first category (constrained loss and gain of targeting) was highly enriched (Fig 4C, 30 miRNAs observed versus 0 expected, binomial P=6.1×10^−58^). We also found many deeply conserved miRNAs that exhibited strong selection against loss of existing target sites, but showed weak or no selection against gain of novel sites (23 miRNAs versus 3.5 expected, binomial P=3.9×10^−13^). We observed a group of deeply conserved miRNAs in the third category, such as miR-146, whose targeting was not conserved beyond the 5% threshold for either loss or gain of target sites.

Consistent with our earlier analysis (Fig 2), this third category of target sites with unconstrained target site loss and gain rates for target sites was depleted (20 observed vs. 67 expected, binomial P=3.1×10^−37^). Only two miRNAs showed constraint against gain without constraint against loss of target sites (which was not significantly different from the expected 3.5). Taken together, these results suggest the presence of three main groups of miRNAs within the deeply conserved miRNAs, with each group exhibiting pronounced differences in the evolution of their target site complement. There are those whose existing target sites are constrained against loss, and which avoid accumulation of novel target sites (group 1), there are those whose existing targeting is constrained, but with unconstrained accumulation of novel targets (group 2), and there are those for which rates of target site loss and gain are not constrained (group 3). The large number of miRNAs in group 3 is somewhat surprising given that these deeply conserved miRNAs are conserved in sequence across the span of vertebrates from human to mouse to zebrafish.

To gain additional perspective on the evolution of animal miRNAs and their target sites, we repeated our analyses on a Drosophila phylogeny including seven species (*melanogaster*, *simulans*, *sechellia*, *yakuba*, *erecta*, *biarmipes*, and *suzukii*; S1 Fig). In this dataset, we defined deeply conserved miRNAs as those present in *D. melanogaster*, *Triboleum castaneum*, and *Apis mellifera* (fly, beetle and bee, respectively; Fig 4D, S9 and S10 Tables), resulting in 43 deeply conserved insect miRNAs. It should be noted that this definition of a ‘deeply conserved’ miRNA is somewhat more stringent than our earlier vertebrate definition, as flies, beetles, and bees are more diverged than humans, mice, and fish (*D. melanogaster*, *T. castaneum*, and *A. mellifera* have incurred a summed branch length of 3.8 substitutions per site since diverging from a common ancestor, whereas *H. sapiens*, *M. musculus*, and *D. rerio* have incurred a summed branch length of 2.6 substitutions per site since diverging from a common ancestor). Our analysis of insect miRNAs therefore contains a smaller set of more deeply conserved miRNAs than we used when examining deeply conserved animal miRNAs. Similar to our mammalian results, we found that deeply conserved insect miRNAs tend to lose and gain targeting at rates significantly below expected rates (S2 Fig) We also found that the three groups of deeply conserved miRNAs we described in mammals also manifest in the Drosophila phylogeny (Fig 4D). Using ranks below 5% of all 8mers to indicate constrained target sites, 24/43 deeply conserved insect miRNAs exhibit reductions in both loss and gain of target sites (group 1, enrichment binomial P=1.9×10^−15^). Nine of the

Drosophila miRNAs were constrained for target site loss and unconstrained for target site gain (group 2, enrichment binomial P=1.6×10^−4^). 9/43 highly conserved miRNAs were not strongly conserved against creation or destruction of binding sites and therefore appeared to be relatively promiscuous in their interactions with targets (group 3, depletion binomial P=9.7×10^−27^), and only one highly conserved miRNA had strong selection against creation of novel sites, but weak selection against loss of existing sites (group 4, binomial P=0.39). Thus, patterns of evolution of miRNA target sites in mammals are recapitulated in Drosophila.

### Older miRNAs are broadly expressed and under stronger selection than younger, more narrowly expressed miRNAs

Following up on our results indicating that deeply conserved miRNAs fall into three groups, based on patterns of target site loss and gain, we next searched for parameters that might explain these differences. We examined miRNA and target expression data to determine whether differences between the efficacy of selection leading to creation or destruction of miRNA target sites correlates with differences in the expression patterns of these miRNAs. Using a miRNA expression atlas [46], we found that miRNAs with strong selection against both loss and gain of targeting (those in group 1) tend to be more broadly expressed (defined as expression at any level) than miRNAs with strong selection against gain of targeting and weak selection against loss of targeting (those in group 2, Wilcoxon Rank P=0.006, bottom of S15 Table). miRNAs with weak selection against both gain of targeting and loss of targeting (in group 3) were generally expressed in fewer tissues than those in group 1 (S15 Table; group 1 vs. group 3 P=0.0004, group 2 vs. group 3 P=0.0526, Wilcoxon rank sum tests). miRNAs that are expressed broadly across the entire organism therefore appear to exhibit greater conservation of targeting than miRNAs that are narrowly expressed. However, we observed numerous exceptions to these trends; for example, miR-146 is broadly expressed and yet exhibits negligible constraints in gain and loss of target sites, while other miRNAs, such as miR-137, are narrowly expressed and yet exhibit strongly constrained patterns of target site loss and gain (S15 Table). We conclude that miRNA expression patterns are correlated with patterns of target site evolution, but there are unambiguous exceptions to the overall correlation, and therefore patterns of miRNA expression and target evolution are not likely to be deterministically linked.

### Differences in patterns of miRNA target site evolution across the transcriptome

An outstanding question in the miRNA field concerns how many meaningful targets exist amongst the hundreds of possible target sites for a given miRNA. Possible answers to this question range from beliefs that each miRNA possesses a small handful of targets whose regulation is consequential, to hypotheses that each miRNA regulates hundreds of targets [47]. Because many miRNAs are highly tissue specific [46], only a fraction of potential targets are coexpressed with their cognate miRNA. To gain additional perspective on this question, we examined the relative strength of selection acting on individual mRNAs for each miRNA. We began by analyzing each individual 3′UTR and comparing the number of observed substitution events against the mean of the simulated datasets for each possible 8mer. Some 3′UTRs incurred far fewer substitutions than expected, while other genes incurred far more. However, even in a randomly generated dataset, some points will fall far below the mean, and others far above. To control for this stochastic fluctuation, we repeated this process multiple times, choosing one of the simulations to compare against the mean of all others (Fig 5, purple lines; S1 zipped file). This process revealed that for many of the deeply conserved miRNAs, the observed selective pressure against loss events is stronger than any of the simulated datasets. For some deeply conserved miRNAs, exemplified by let-7, we observed reduced rates of target site loss substitutions and reduced rates of target site gain substitutions across a large fraction of the 3′UTRs, indicating that almost all of these targets or potential targets are under selection (Fig 5A-B losses and gains, left and right, respectively). In contrast, other deeply conserved miRNAs, exemplified by miR-21, showed evidence of negative selection only for a small subset of the target space, suggesting functions that are restricted to a small portion of the possible set of regulated transcripts (Fig 5C-D, losses and gains, left and right, respectively), with the majority of the potential target set evolving at near neutral rates. Thus, the patterns of target site evolution for some miRNAs are consistent with broad selection acting across many targets, whereas for other miRNAs, selection is most apparent on a more restricted target set. Some miRNAs, exemplified by miR-146, showed very weak, if any, evidence of selection across most of the transcriptome (Fig 5E-F, losses and gains, left and right, respectively), while still other miRNAs showed subsets of highly constrained targets in addition to subsets of targets with weak evidence of rapid turnover of binding sites (Fig 5G-H, losses and gains, left and right, respectively). In this manner, our method demonstrates an ability to differentiate deeply conserved miRNAs that appear to operate on large subsets of genes from those that operate on small subsets of genes, and to differentiate between miRNAs that operate only in conserved processes, those that operate in both conserved and rapidly evolving processes, and those that appear to operate predominantly in rapidly evolving processes. For example, certain ancient miRNAs, including let-7, miR-9, and miR-1, exert strong purifying selection on target sites across much of the transcriptome (Fig 5, S1 zipped file), while other ancient miRNAs, including miR-21, appear to only exert purifying selection on only a small subset of the transcriptome; in contrast, we also identified evidence of rapidly changing targets for conserved miRNAs such as miR-146. By analyzing the distribution of selection across genes, our approach is able to differentiate strong selection acting on a small number of target mRNAs from weak selection acting broadly across the transcriptome, facilitating a deeper understanding of the types of roles that individual deeply conserved miRNAs are likely to play.

**Fig 5.**
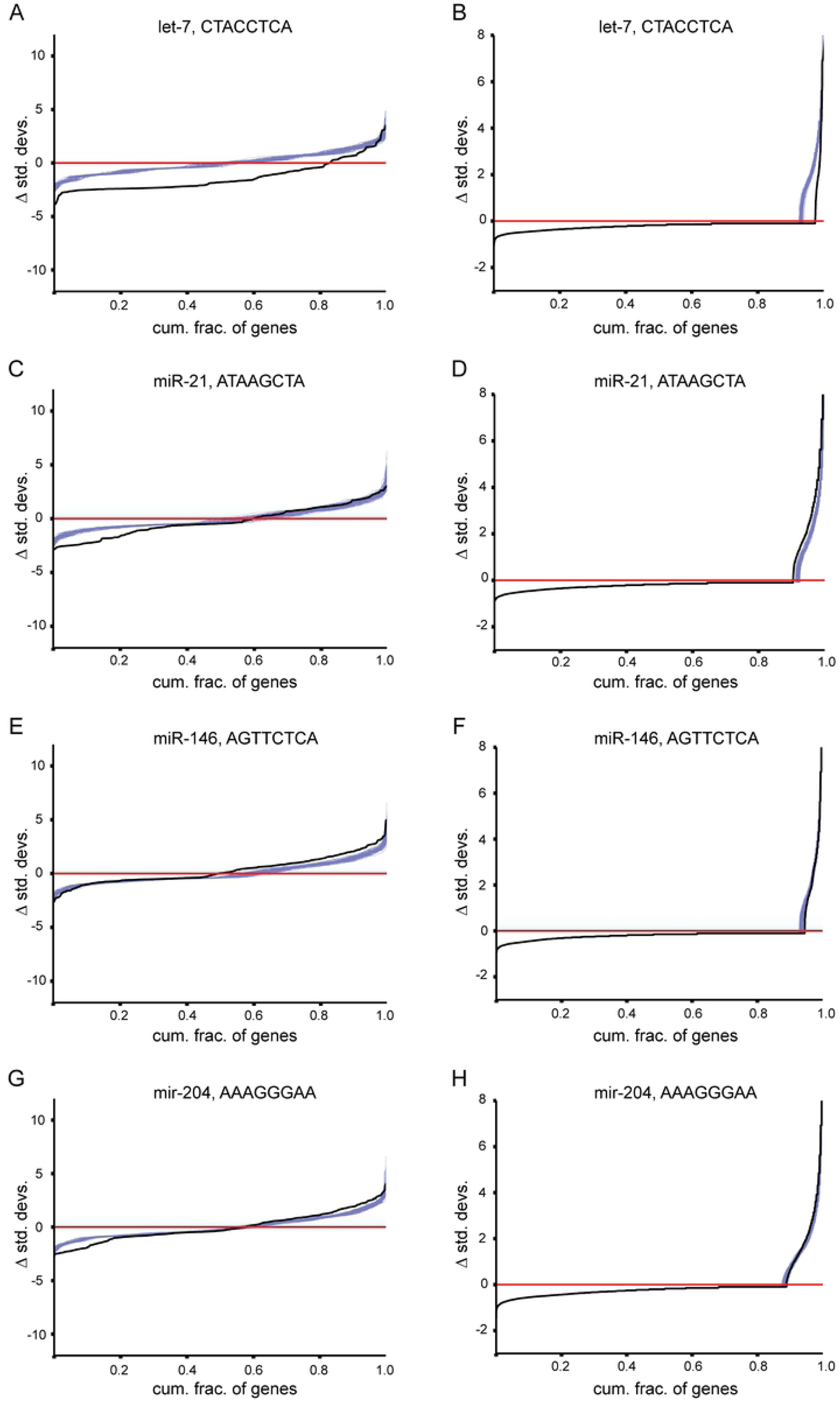
Target sites for individual miRNAs exhibit distinct patterns of selection. For each miRNA, all 3′UTRs that undergo substitutions that destroy or create an 8mer miRNA target site were analyzed. For each 3′UTR, the total number of such substitutions was evaluated relative to the mean of simulated substitutions for that 3′UTR, and the number of standard deviations above or below the mean recorded (y-axis), ordering genes from those that fall the greatest number of standard deviations below the mean to those that fall the greatest number of standard deviations above the mean (x-axis). The black line represents scores for the observed dataset as compared to the mean of 99 simulated datasets. The purple lines represent scores for each of the 100 simulated datasets as compared to the mean of the remaining 99 simulations, and serve as a measure of expected stochastic fluctuations. Regions in which the black line is above all other lines represent an excess of substitution events (positive selection favoring substitutions), while regions in which the black line is below all other lines represent a depletion of substitution events (negative selection against substitutions). **(A, C, E, G)** Analysis of target site loss rates (left panels) for the 8mer target sites corresponding to the miRNAs let-7, mir-21, mir-146, and mir-204, respectively. **(B, D, F, H)** Analysis of target site gain rates (right panels), otherwise as for panels A, C E and G.

### Changes in miRNA target site efficacy are constrained

MicroRNA target sites exist with a range of efficacies. In general, the weakest sites are those with a 6 nucleotide match (6mers), created by pairing of miRNA nucleotides 2-7 with 3′UTR sequence. The most effective sites (8mers) supplement the core 6 nucleotide match with both an additional base pair to position 8 of the miRNA (m8) and an unpaired A (A1). We therefore focused primarily on 8mer sites, reasoning that signatures of selection would be most pronounced for the strongest class of target sites. There are also seven nucleotide target sites, typically intermediate in efficacy between 6mer and 8mer sites, which contain either the m8 or A1 nucleotide; 7mer-m8 sites tend to be, on average, more effective than 7mer-A1 sites [19]. Although the existence and relative efficacies of these different classes of miRNA target sites are well established, the relative strength of selection operating to convert site-types from one state to another has not been examined previously.

We used our simulated 3′UTR datasets as a benchmark to search for constraints on evolutionary changes between the four classes of miRNA target sites, examining those corresponding to the deeply conserved miRNAs only. We compared the frequency with which target sites convert between site types to the frequency with which other, background, 8mers convert between site types, in order to obtain estimates of the strength of selection operating on these events. We found that miRNA target site conversion events of almost every type tend to be significantly depleted when compared to equivalent events acting on background 8mers, which we assumed approximate neutral evolution (Fig 6, SFig 3). Nearly every target site type demonstrates selective pressure against both strengthening existing binding sites (Fig 6A, beneath and to the left of the shaded diagonal) and against weakening existing binding sites (above and right of the shaded diagonal). Target sites of miRNAs in group 1 (those with constrained gain and loss rates; Fig 4) show particularly strong constraint against conversions (Fig 6B), miRNAs in group 2 (constrained against losses but not gains) show evidence of constraint against weakening target sites and weaker evidence of constraint against strengthening target sites (Fig 6C), while target sites for miRNAs in group 3 (whose 8mer target sites are evolving neutrally) are changing at rates close to neutrality, as expected (Fig 6D). These results suggest that modifications to the strength of an existing miRNA target site are sufficiently consequential to impact the evolution of 3′UTR sequence, and that weakening of an existing site is generally more harmful to the organism than strengthening one (blocks are darker in upper right diagonals than lower left, Wilcoxon Rank p-value 5.6×10^−9^). Because selection appears to act not only against creating novel deleterious sites but also against strengthening existing beneficial sites, we note that these results are most consistent with the hypothesis that many miRNA target sites exist to modulate or fine-tune gene expression rather than to maximally repress gene expression.

**Fig 6.**
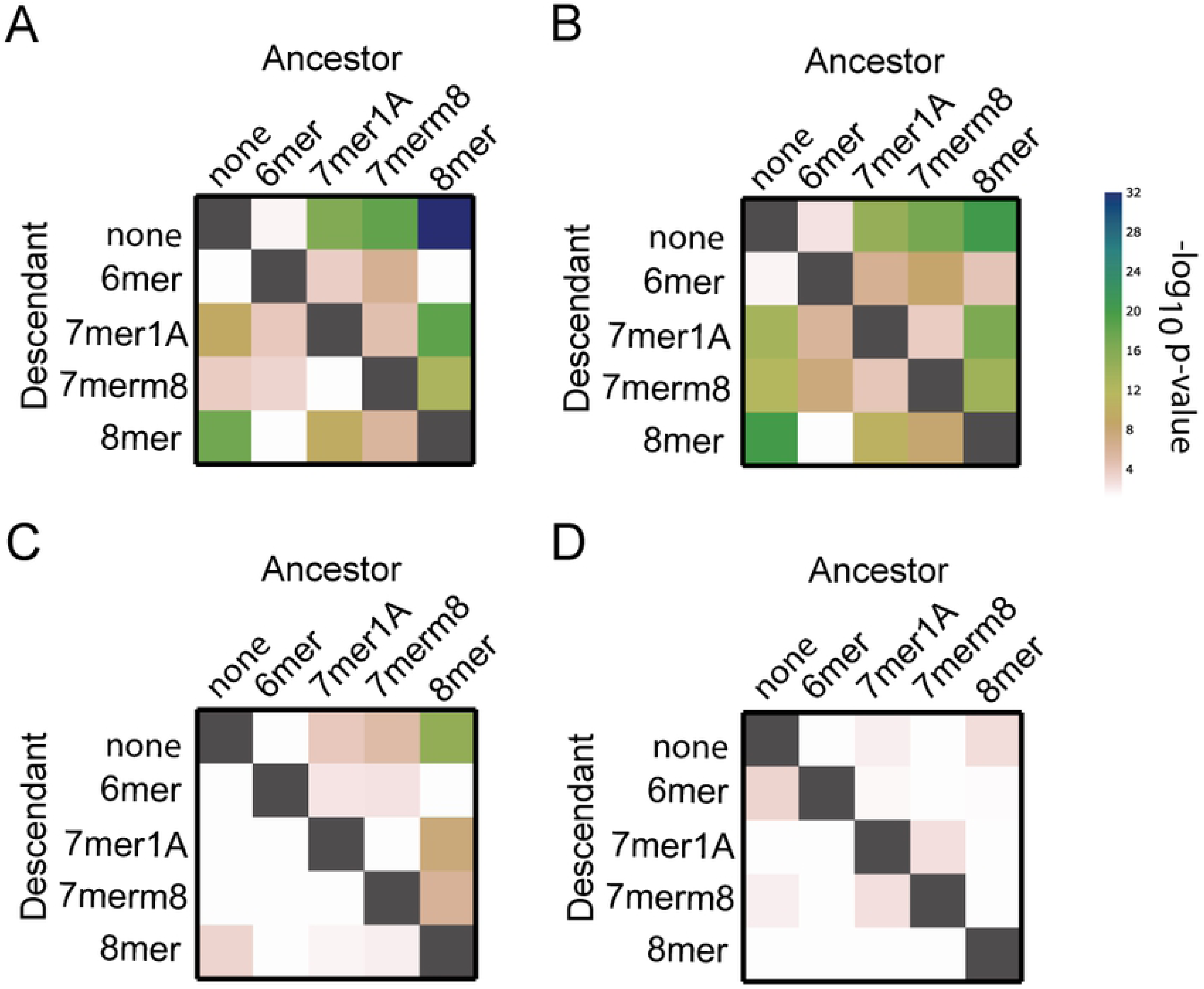
Interconversions that strengthen or weaken miRNA target sites are disfavored. Each panel calculates pairwise comparisons between the foreground and background rates at which the indicated miRNA target sites undergo substitutions (see methods). The ancestral target site type is represented along the x-axis, and the descendant along the y-axis. Substitutions that increase the average strength of a target site are below and to the left of the diagonal (grey boxes), while substitutions that decrease the average strength are above and to the right. Significance is color-coded (see legend) and plotted as -log_10_(P-value), using Wilcoxon rank-sum comparisons between the background and foreground rates. Insignificant P-values (P>0.05) are color coded white. **(A)** Analysis of target site conversion events for all deeply conserved miRNAs **(B, C, D)** Analysis of target site conversion events for deeply conserved miRNAs with strong selection against both gain and loss events (B; referred to as group 1 in main text), miRNAs with strong selection against loss events but not gain events (C; group 2), and miRNAs with no strong selection against loss or gain events (D, group 3).

### Analysis of most constrained 3′UTR 8mer motifs

The regulatory information within 3′UTRs extends beyond that mediated by miRNAs [48]. Therefore, we decided to systematically examine all 8mers for selective constraint in mutations that either create or destroy the 8mer. We examined the 1,000 most constrained 8mers (1.5% of all 8mers), ranked independently for losses and gains. As expected, 8mers that correspond to or overlap with miRNA target sites represent a large fraction of the most constrained 8mers. We also examined 8mers with overlap to Pumilio binding sites, cleavage and polyadenylation signals, AU-rich elements and Fox binding sites [49–51]. Cumulatively, these established 3′UTR regulatory elements represent 66.8% and 48.1%, respectively, of the 8mers with most constrained losses and gains. As a background set, we repeated this analysis on 1,000 unconstrained 8mers (Fig 7), which had only 13.9% and 13.2% of elements deriving from known motifs for losses and gains, respectively. Thus, binding sites for established 3′UTR regulatory elements exhibit marked constraints in sequence changes that either create or destroy existing sites. It seems likely, therefore, that our approach could be used to identify novel regulatory 3′UTR motifs by searching for sequence motifs that are both rarely lost and gained during evolution.

**Fig 7.**
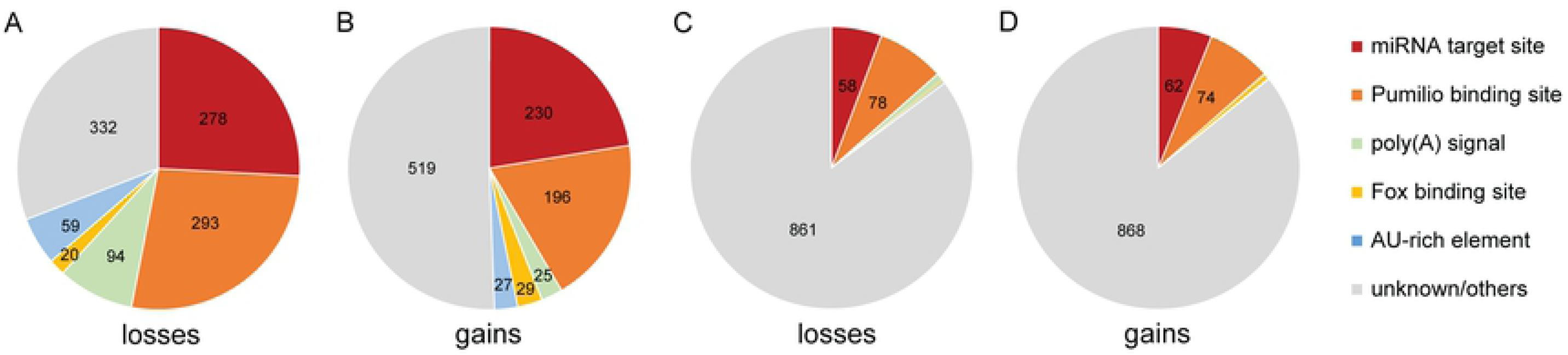
Analysis of rates of loss and gain of functional 3′UTR regulatory elements. **(A, B)** Fractions of the 1,000 most constrained 8mers corresponding to miRNA target sites, Pumilio response elements, poly(A) signal sequences, Fox protein binding sites, AU rich elements, and motifs of unknown function, for losses (A) and gains (B), analyzed as described in Fig 1. **(C, D)** Fractions of the 1,000 least constrained 8mers, otherwise as described in panels A and B.

## MATERIALS AND METHODS

### 3′UTR Sequences

Transcript coordinates were downloaded from the Refseq track of the UCSC genome browser [5]. Transcripts with overlapping 3′UTRs were consolidated, retaining the longest 3′UTR isoform. For analysis using mammalian sequences, we used multiz 100 way alignments [52] to the hg19 reference, examining the ten mammalian species human, gibbon, marmoset, bushbaby, dog, cat, cow, horse, sheep, and mouse. For analysis of *Drosophila* sequences, we used multiz 27 way alignments [52] to the dm6 reference, examining the seven *Drosophila* species *D. melanogaster*, *D. simulans*, *D. sechellia*, *D. yakuba*, *D. erecta*, *D. biarmipes*, and *D. suzukii*. Insertions and deletions of multiple nucleotides were collapsed to a single ‘N’ in all species. 3′UTRs with a collapsed length of less than 50 nucleotides were removed from analysis, as were 3′UTRs in which any species was completely missing from the alignment. The complete datasets comprised 13,045 and 8,755 aligned 3′UTRs for the mammalian and Drosophila phylogenies, respectively.

### Ancestral reconstruction and simulation

We used the aligned 3′UTRs in concert with the phylogeny provided with the UCSC multiz alignments of the dm6 and hg19 genomes as input for the program DNAML, which can be obtained as part of the phylip package [41]. The program outputs inferred ancestral sequences at every internal node of the phylogeny, for every alignment. Every ancestral sequence was paired with each of its descendants, such that every branch of the phylogeny was represented once, and for every ancestor/descendant pair, we searched all trinucleotides within the alignments from all 3′UTR transcripts, in order to capture the effects of immediately adjacent nucleotides on the probability of a substitution. For each trinucleotide, we compared the ancestral trinucleotide to that of the aligned descendant. Because we were interested in the mutation probability of a nucleotide given its two flanking nucleotides, and to avoid double counting mutations, we ignored all trinucleotides in which the first or last nucleotide differed between the ancestor and descendant. Each ancestor/descendant pair thus produced a table of nucleotide substitution frequencies given fixed flanking nucleotides on either side. Because trinucleotides without a mutated central nucleotide were also recorded in the table, both the probability of incurring a substitution and the frequency of each type of substitution were accurately reproduced in the table. We generated one table (aggregated across all aligned 3′UTRs) per branch of the phylogeny.

To generate our simulated dataset, we scanned all ancestral trinucleotides a second time. We used the substitution probability table generated in the first pass to generate a weighted random process of choosing a descendant trinucleotide for each ancestor, and appended the central nucleotide of this randomly generated trinucleotide to a growing simulated descendant sequence. This process was used to generate simulated descendant sequences for all 3′UTRs in all ancestor/descendant pairs, resulting in a complete phylogeny of simulated descendants for all sequences. This process was repeated 100 times to generate 100 simulated descendant 3′UTR datasets in mammals, each containing 13,045 genes in 17 ancestor/descendant pairs. In insects, this process was repeated 100 times to generate 100 simulated descendant 3UTR datasets, each containing 8,755 genes, in five ancestor/descendant pairs.

### Statistical analysis and ranking of expected and observed target site turnover rates

We examined every 8mer in both the simulated and observed datasets, and recorded instances in which the ancestral 8mer differed from its aligned descendant. These events were considered as a ‘gain’ of the descendant 8mer and as a ‘loss’ of the ancestral 8mer. After scanning every ancestor/descendant paired alignment in this way, the result was tallied for each 8mer as a summed count of gain events within the entire dataset (across all branches and genes) and a summed count of loss events within the entire dataset. Because there were 100 simulated datasets and one observed dataset, our final summary recorded one observed ‘gain’ count for each 8mer and 100 simulated ‘gain’ counts, as well as similarly calculated observed and simulated ‘loss’ counts. We calculated the mean and standard deviation of the 100 simulated counts for each 8mer, and calculated the number of standard deviations each observed 8mer count fell above or below the mean of the simulated counts. Finally, the standard deviation calculations for each 8mer were ranked relative to each other in the ‘gain’ and ‘loss’ categories. The strength of selection acting on the mutation rates of an 8mer relative to other 8mers can be approximated from its rank, while a conservative estimate of the probability of observing a similar or more extreme standard deviation was approximated using Chebyshev’s inequality [53], which states that for any distribution of values, the upper bound on the probability of observing a value that is k standard deviations from the mean is P<1/k^2^.

Because our aligned intronic sequence (selecting only regions with no insertions or deletions in any species) was roughly 23 times the size of our 3′UTR alignment, and in order to avoid altering our power to detect selection and avoid intronic splice sites that occur at the outer edges of introns, we examined the central 1/23rd of every intron. Analysis was then performed as above.

### Gene-specific analysis

We began by re-analyzing our dataset using the same techniques as before, but, instead of recording the total number of substitutions that occur across all 3′UTRs, we recorded the total number of substitutions that occurred for each 8mer within each gene. As previously, we compared the observed number of substitutions to the mean and standard deviation of 100 simulated substitutions, in order to count the number of standard deviations that the observed value fell above or below the mean expected value. We plotted this number of standard deviations for each gene on the y-axis, and ordered genes from those that were changing many standard deviations below expected rates to those changing many standard deviations above expected rates. Stochastic processes followed patterns that were not always intuitive. For example, an 8mer that almost never undergoes substitutions will appear to be changing at an extremely rapid rate within the small subset of genes that do undergo substitutions, even if no selection is present. To establish expected rates of substitution, we compared our observed dataset to the mean of 99 simulations, and additionally compared a random simulated dataset to the mean of the other 99 simulations. We repeated this process using every neutral simulation as foreground to generate 100 ‘neutral foreground’ datasets and one ‘selected foreground’ dataset, and plotted the results (S1 zipped file).

### Removal of miRNA target sites perfectly conserved across analyzed 3′UTRs

We first located all positions in our observed alignments that have undergone substitutions in at least one descendant species, creating a set of mutated sites. We next counted the number of transition events (gains or losses) that occurred for 8mers overlapping by at least one nucleotide with these mutated sites. This resulted in an elimination of all perfectly conserved 8mer sites from the data. To ensure equally sized and identical pools of simulated ancestral 8mers to the observed dataset, we also examined the same coordinates within simulated alignments (regardless of whether a substitution occurred at these locations). Thus, our observed dataset contained only ancestral 8mers that have undergone at least one substitution in a descendant species, while our simulated dataset contained the same ancestral 8mers, regardless of whether they have undergone a substitution. The resulting observed and simulated datasets were used to examine whether miRNA binding sites that have undergone at least one substitution are constrained relative to other 8mers that have undergone at least one substitution.

### miRNA site-conversion rates

miRNA 8mer target sites can be conceived as a 6mer ‘core’ site, which basepairs to nucleotides 2-7 of the miRNA, with two specific additional flanking nucleotides. Each flanking nucleotide alone converts a 6mer site into a different type of 7mer site. An 8mer→7mer_1A substitution is caused when the target site undergoes a substitution at its first nucleotide, an 8mer→7mer_m8 substitution is a substitution at the last nucleotide, an 8mer→no site substitution is any substitution in the core 6 nucleotides, with equivalent conversions calculated between other classes of target sites. These principles were generalized for all 8mers. By examining rates at which all 8mers undergo mutations at their first nucleotides, we can compare rates of 7mer_m8→8mer and 8mer→7mer_m8 conversions for both miRNAs and other 8mers, which were used as controls. We repeated this analysis for all site conversion types (6mer→7mer_m8, 6mer→7mer_1A, 6mer→8mer, none→8mer, 7mer_m8→8mer, 7mer_1A→8mer, none→6mer, none→7mer_1A, none→7mer_m8 constituted our ‘gain’ event types, while each of these in reverse constituted site ‘loss’ events). Each 8mer conversion rate was ranked relative to the same conversion rate for all other 8mers as in our initial analysis, and as before, we performed a Wilcoxon rank-sum test to evaluate whether the set of target sites of conserved miRNAs have ranks that tend to fall significantly above or below ranks of the complete set of all 8mers.

## DISCUSSION

We have set out here to undertake a comprehensive analysis of the evolutionary relationships between miRNAs and their targets. Previous work has demonstrated that conservation of a target site can be used to predict functional sites [21, 33, 34, 54, 55]. The existence of ‘antitargets’ of miRNAs, that is, 3′UTRs under selection to avoid containing miRNA target sites, has also indicated that selection acts to prevent the gain of detrimental target sites [30, 56]. Many additional studies have made valuable contributions to our understanding of the origins and evolution of miRNA target sites [20, 31, 38, 42, 57]. However, a systematic evolutionary analysis of all changes, that is, rates of gains and losses in individual target sites of individual miRNAs, constitutes an important and largely unexamined component of miRNA biology. The role of a miRNA is defined by the collection of genes that it targets. Thus, it is only by systematically studying changes in targeting that we can gain insights into the ways in which individual miRNAs change roles over deep evolutionary time. Our goal here was to construct a reliable background model of 3′UTR sequence change, and use this model to infer rates of miRNA target site gain and loss attributable to selection. This approach has allowed us to gain new insights into the changing roles of miRNAs during evolution, including at the level of individual miRNAs.

We find strong selection against loss of the target sites of deeply conserved miRNAs (Fig 2A), consistent with previous studies [34]. We find comparably strong selection against target site gain (Fig 2B); although this result has been hinted at previously [30], the extent of the signal is surprising, and comparable to the selection against losses. While deeply conserved miRNAs might be expected to have acquired deeply conserved targets, the effects of miRNAs on gene expression are subtle, and the creation of accidental target sites would typically exert relatively modest effects on gene expression levels. The broad extent of the signal against target site gain indicates that for most deeply conserved miRNAs, even subtle reductions in gene expression across potential targets are strongly selected against. Importantly, when we repeated these analyses in Drosophila species, we observed strong selection against both gain and loss of miRNA target sites (S2 Fig), suggesting that these patterns of selection acting on miRNA target site gain and loss are likely to represent diverse bilaterian lineages.

### Patterns of target site loss and gain for individual miRNAs

In general, miRNAs with the slowest turnover rates of target sites are in good agreement with those described as having ancient functions. For example, the let-7 miRNA is known to have at least one target, lin-41, which has a target site that is maintained in organisms from nematodes to humans [31], and our analyses indicate that let-7 is highly constrained, both in terms of target site loss and gain across mammals. Similarly, the target sites for other well-studied and ancient miRNAs, such as miR-124 and miR-9, are also highly constrained. Interestingly, the two miRNAs that stand out as having the slowest rates of change in targeting, miR-20 and miR-30 (S1 and S2 Tables) are not prominently described in the literature as having deeply conserved targeting. Both miR-20 and miR-30 are expressed in a wide range of tissues, and are amongst the most broadly expressed miRNAs (S15 Table). The success of our method in enriching for deeply conserved miRNAs with known deeply conserved functions implies that miR-20 and miR-30 also have deeply conserved functions. miR-30 has been shown to play a role in thermogenesis, which may form part of a conserved pathway among mammals, and appears to play important roles in adipogenesis and intestinal epithelial cell homeostasis [58–60], while miR-20 has roles in suppressing angiogenesis and is a member of a miRNA family that is essential in mice [61–63]. Thus, both of these miRNAs may represent promising candidates for the discovery of conserved and essential miRNA-mediated regulatory pathways among mammals, and potentially other bilaterian species.

In contrast to let-7 and many other ancient miRNAs, miR-146 is an unusual example of a deeply conserved miRNA whose target set is minimally constrained. Indeed, miR-146 target site losses and gains are observed to occur at rates exceeded by only ∼10% of all 65,536 surveyed 8mers. These observations indicate that this miRNA is gaining novel targets and losing existing targets extremely rapidly, with neutral or potentially faster than neutral rates. Notably, miR-146 contributes to diverse immune system functions, including fever response, tumor suppression, T-cell homeostasis, and cytokine production in multiple immune cell lineages [64–68]. It is conceivable that miR-146 targets may be rapidly co-evolving with pathogens as part of an ‘arms race’ scenario that has frequently been seen in host-pathogen interactions [69]. Thus, miR-146 may represent a promising candidate for the discovery of novel rapidly evolving (non-constrained) regulatory pathways that harness specific deeply conserved miRNAs for these purposes.

We also considered miRNAs beyond those we identified as deeply conserved, none of which exhibited evidence of strong selection against site loss or gain (Fig 2), consistent with previous studies indicating a lack of conservation of target sites for such miRNAs [42]. The existence of so many miRNAs, including many that are broadly conserved across mammals, which exhibit minimal or no evidence of selective pressure on their target sites remains somewhat perplexing. The simplest interpretation of these results is that such miRNAs have a very restricted set of targets that are under selection, but that such targets are greatly outnumbered by those that are evolving at neutral rates. Alternative possibilities, which have already been established in the evolution of transcription factor binding site (TFBS) networks [8–10, 16] and posited for miRNA target site networks [35], include the existence of a set of targets that are evolving rapidly, but that are biologically consequential. In this regard, it is important to note that we detected selection against target site loss and gain for deeply conserved miRNAs even when we restricted our analysis to regions of 3′UTR sequence undergoing more rapid sequence change (Fig 3). This result indicates that even when only weakly conserved target sites are analyzed, deeply conserved miRNAs still exert significant selective pressure against sequence change, while less deeply conserved miRNAs show no detectable selection against losses or gains of target sites, even when more deeply conserved 3′UTR sequences are included. Thus, over a relatively short evolutionary timescale, selection acts on targets corresponding to ancient miRNAs, but does not act on less well-conserved miRNAs, even though such miRNAs pre-date the timescale considered. It is clear that the evolutionary ages of miRNAs plays a larger role in the strength of selection acting across the transcriptome than the age of individual target sites.

### MiRNAs can be classified into distinct groups

We note the existence of three markedly different groups of deeply conserved miRNAs, each with characteristic patterns of target site evolution. The first group contains miRNAs whose target sites are constrained for both losses and gains, and is the largest of the three (Fig 4C, D). MicroRNAs of this type are to be expected, as it is clear that selective pressures exist that maintain existing sites [34] and preclude the evolution of new sites [30], which would lead to reductions in losses and gains, respectively. The second group contains miRNAs which exhibit gains of sites at neutral rates but that are still constrained against loss of existing sites (Fig 4C, D). Such miRNAs are more likely to gain new consequential sites across mammalian evolution than the first group, for which gain of new sites is selected against and comparably rare. The existence of this second, large, group of miRNAs is unexpected, as it indicates that some miRNAs have deeply conserved existing targets, yet gain novel targets at approximately neutral rates. The patterns of target site evolution of the third and final group, which includes many ancient miRNAs, are the most surprising, in that they do not exhibit constraints on target site loss or gain. While this third group has parallels in transcription factor networks that are known to include deeply conserved transcription factors with rapidly shifting targets [16], this group is largely unanticipated within the miRNA literature, and represents miRNAs that appear to be deeply conserved while gaining and losing target sites at approximately neutral rates. It is important to acknowledge the existence of this final class, as it represents miRNAs for which conservation of individual target sites is less likely to represent a meaningful signal. In addition, the existence of this class of deeply conserved miRNAs presents a potential mechanism that would allow deeply conserved miRNAs to drive striking shifts in gene regulatory networks in a small amount of time without any change in miRNA sequence. Amongst the most deeply conserved miRNAs, only two (of 75) exhibited selection against site gain while losing sites at a neutral rate. Thus, our results indicate that it is almost never the case that a deeply conserved miRNA exhibits strong deleterious effects as novel target sites accumulate when no selection is detectable against loss of existing targets. By classifying miRNAs into these groups, it is now possible to make reasonable conjectures as to which deeply conserved miRNAs are most likely to be involved in deeply conserved processes, which are expanding into novel functional niches, and which might potentially be associated with rapidly evolving species-specific functions.

The number of consequential targets of individual miRNAs has been a subject of some debate. While preferential conservation of miRNA target sites within 3′UTRs demonstrates that hundreds of genes are evolving in response to individual miRNAs [34, 42], others have shown that the phenotypes resulting from loss of function of miRNAs can often be recapitulated or amerliorated by altering the experession of a very small subset of target genes [70, 71]. Moreover, the narrow expression patterns observed for some miRNAs suggests highly tissue-specific functions [72]. Still other studies posit that miRNAs function partly to repress “leaky” or “noisy” gene expression of a large number of genes whose expression is undesirable in specific cells or a stages of development [73, 74]. Under this model, even highly tissue-specific miRNAs may still consequentially regulate large cohorts of targets as a consequence of selection; it is important to note that such targeting may be difficult to robustly test or detect using experimental methods. Our method goes beyond traditional metrics of sequence conservation by examining individual targets of miRNAs for evidence of selective constraint against target site turnover (Fig 5). We have observed that transcriptome-wide or gene specific selection on target sites is not a binary trait that can be applied homogeneously to all deeply conserved miRNAs. While changes in target sites corresponding to some miRNAs exhibit selective pressure across much of the transcriptome, others exhibit strong selective pressure on a very small number of targets. Our approach may therefore be applied in novel ways to distinguish between miRNAs that serve as broad-spectrum regulators of the genome and those with very specific, restricted functions. Importantly, our results imply that different deeply conserved miRNAs have evolved under markedly different modes of selection, and suggest that no single theory of miRNA targeting is likely to be broadly applicable.

### Identifying and characterizing selective pressures acting on novel motifs

Recent studies by multiple groups, using a variety of approaches, indicate that regulation by miRNAs is only a modest fraction of the regulatory information found in mammalian 3′UTRs [75–79]. While our method has been applied primarily to the characterization of selective pressures acting on miRNAs, we also analyzed selective pressures operating on all possible 8mer motifs, and successfully recovered motifs corresponding to RNA binding proteins with regulatory functions. We identified AU-rich elements, Fox and Pumilio binding sites, and polyadenylation signal sequences (Fig 7) as motifs with slow rates of loss and gain. Our approach is therefore capable of characterizing selective pressures acting on a wide range of nucleotide sequence motifs. Importantly, we also found a large number of sequences with slow rates of loss and gain in 3′UTRs that do not correspond to previously recognized regulatory elements, such motifs may represent novel 3′UTR functional elements.

### Fine-tuning by MicroRNA target sites

A diversity of functional roles have been proposed for miRNA target sites. Broadly speaking, these roles constitute three classes: subtle quantitative control of gene expression, often referred to as fine-tuning [47], miRNA-mediated switches that seek to maximally repress or completely turn off a gene, or contribute to this type of regulation in concert with other mechanisms [47], and sites that titrate miRNAs to a level optimal for regulation of a very small subset of true targets, an idea encapsulated by the competing endogenous RNA (ceRNA) hypothesis, [80, 81]. Undoubtedly all three classes exist, but their relative prevalence and importance is less certain. Our data indicate that substitutions that either strengthen or weaken existing target sites are disfavored (Fig 6); the simplest interpretation of this result is that target sites exist predominantly to optimize levels of the targeted transcript. In contrast, if miRNA target sites acted typically to maximize repression of gene expression, we would expect mutations that increase site efficacy to be selected for; aggregated across miRNAs, we observe the opposite: selection against sites increasing in strength. The ceRNA hypothesis implies selection maintaining the total number and strength of sites with minimal selection acting on their location across the transcriptome: our results are directly opposed to this expectation in that we observe selective pressure acting against the creation and destruction of most target sites, consistent with biochemical studies that imply ceRNAs are exceedingly rare [82, 83]. Thus, our results are most consistent with the model that miRNAs often serve to precisely modulate, or fine-tune, the levels of their target transcripts, as evidenced by selection not only maintaining the locations and numbers of sites, but the potency of the sites themselves.

### A model of miRNA aging

We observe that many miRNAs known to have deeply conserved targeting, such as miR-9 and let-7, and additional miRNAs that we identified, such as miR-20 and miR-30, are also broadly expressed. In general, we find that miRNAs with the strongest constraint against gain and loss of target sites (those in group 1) tend to have broader expression patterns than miRNAs with weaker constraint against gain of targeting (those in group 2, S15 Table) and much broader expression patterns than miRNAs with weak constraint against gain and loss of targeting (those in group 3, S15 Table). However, it is important to note that there are also exceptions. Among miRNAs that defy expected relationships between expression and conservation of targeting, miR-137 is among the most narrowly expressed in the dataset (S15 Table), and yet experiences stronger combined genome-wide selection against the acquisition of novel target sites and the destruction of existing sites than all but 4 miRNAs (miR-20, miR-30, let-7, and miR-9; S1 and S2 Tables). Furthermore, when we examined gene-specific selection for miR-137, we observed broadselection across the transcriptome against both loss and gain of targeting (see S1 zipped file for the miR-137 seed ‘AGCAATAA’). At the other extreme, miR-146, which gains and loses miRNA targets faster than any other deeply conserved miRNA, is broadly expressed, being detected in most tissues. These results illustrate that while there is a trend for broadly expressed miRNAs to show more constrained evolution of targeting than narrowly expressed miRNAs, it is quite possible for a narrowly expressed miRNA to have highly constrained evolution of targeting, and, for a broadly expressed miRNA to gain and lose target sites rapidly.

In addition to a correlation between conservation of miRNA targeting (as assessed by miRNA group) and miRNA expression (bottom portion of S15 Table), we also observe that miRNAs in group 1 are conserved across a longer total branch length as regulators than miRNAs in group 2 (though not significantly so), and that miRNAs in group 1 are conserved across a significantly longer total branch length than miRNAs in group 3 (bottom portion of S15 Table). Combining these observations with the broad expression patterns of miRNAs having the slowest gain rates, it seems feasible that in general, as miRNAs age and are expressed more broadly, this process allows novel targets of these miRNAs to become increasingly likely to be exposed to negative selection in at least one tissue, effectively ‘trapping’ miRNAs with the strongest deleterious effects as novel target sites arise into regulating the targets they had already established, and hindering them from accumulating more. Under this model, a miRNA that originates with strong selection against novel targets and weak conservation of existing targets is likely to lose its existing targets without acquiring new beneficial ones. Such a miRNA would quickly lose its function and be unlikely to become deeply conserved. This model provides a ready explanation for the dearth of miRNAs that have strong selection against gaining new targets and weak selection against losing existing targets (the leftmost rectangles of Fig 4 C, D). Among miRNAs with strongly deleterious effects for novel targets, only those with highly beneficial existing targets (and therefore strong selection against losing targets) are likely to be preserved. Meanwhile, miRNAs with narrow expression profiles, or other causes of weak selection against novel targets [84], are capable of exploring novel targeting interactions, and can be conserved so long as a meaningful fraction of existing targets are beneficial. Our results also highlight notable exceptions to this trend that may represent fruitful topics for further study. For example, miR-191 and miR-137 have target sites that are gained and lost very rarely despite being younger than almost all deeply conserved miRNAs, while miR-338 and miR-203a have target sites that are gained and lost relatively rapidly despite being older most miRNAs. However, the overall trend toward miRNAs with more constrained target site evolution tending to be older (bottom portion of S15 Table), suggests a model of miRNA aging that is consistent with earlier work indicating that miRNAs generally acquire targets as they age [38]. This classification may guide the search for essential, deeply conserved functions of relatively understudied miRNAs of the first class, like miR-30 and miR-20, while suggesting clade or even species-specific roles for deeply conserved miRNAs of the third class, like miR-146.

### Overall significance

We have demonstrated the utility of our technique for examining a wide range of evolutionary questions. We have examined the relative selective pressures operating on both deeply conserved and more recently evolved miRNAs. We have shown that for deeply conserved miRNAs, even target sites that have been completely lost in at least a subset of mammalian species still undergo substitutions at a rate that is lower than other comparable 8mers. We have presented evidence that substitutions that strengthen or weaken existing miRNA targeting are disfavored. A systematic examination of the rates at which miRNA target sites undergo substitutions makes these questions tractable. In addition, by quantifying rates of sequence motif gain and loss, our approach holds promise as a general method for studying regulatory sequence elements (Fig 7).

A thorough examination of the dynamics that have existed between miRNAs and their targets over time gives important information about the coevolution of miRNAs and their target genes, a subject that we are only beginning to understand. Our approach successfully recovers a signal for the binding sites of deeply conserved miRNAs, and has direct applications for improved miRNA functional analysis. Taken together, our approach has the ability to identify the overall strength of selection acting on both the loss and gain of miRNA binding sites, as well as alterations in strength of targeting and the distribution of natural selection across the genome. By correlating these metrics with the age and expression pattern of a given miRNA, we hope to shed light on the varied and changing roles of individual deeply conserved miRNAs.

## ACKNOWLEDGMENTS

This work was supported by R01GM105668 from NIH to A.G. We thank Emily Davenport and Giles Hooker for helpful discussions.

---

**S1 Fig.**
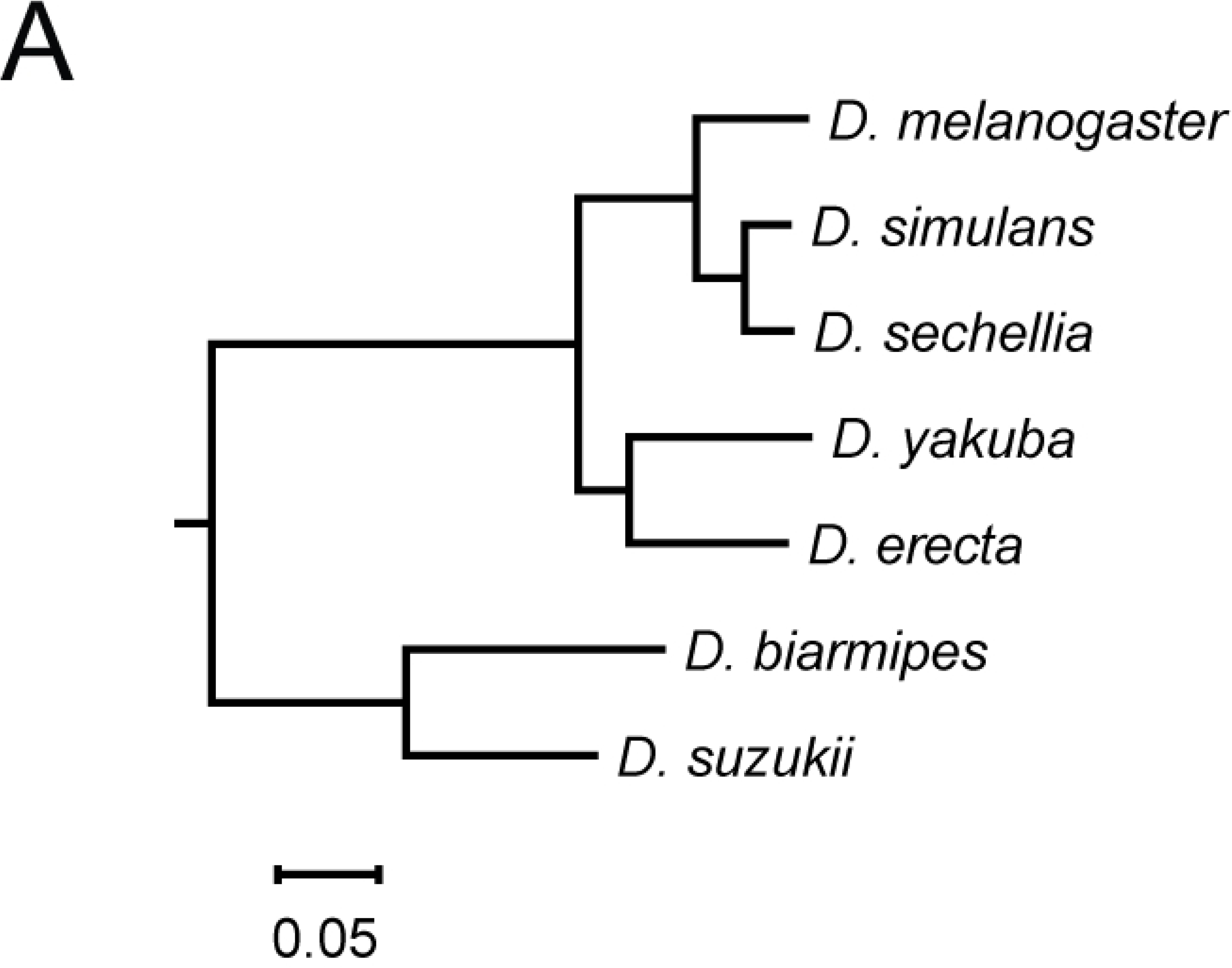
Drosophila phylogeny. Relative phylogenetic distances between selected insect species, measured in substitutions per site, derived from the UCSC genome browser’s multiz 27way insect alignment.

**S2 Fig.**
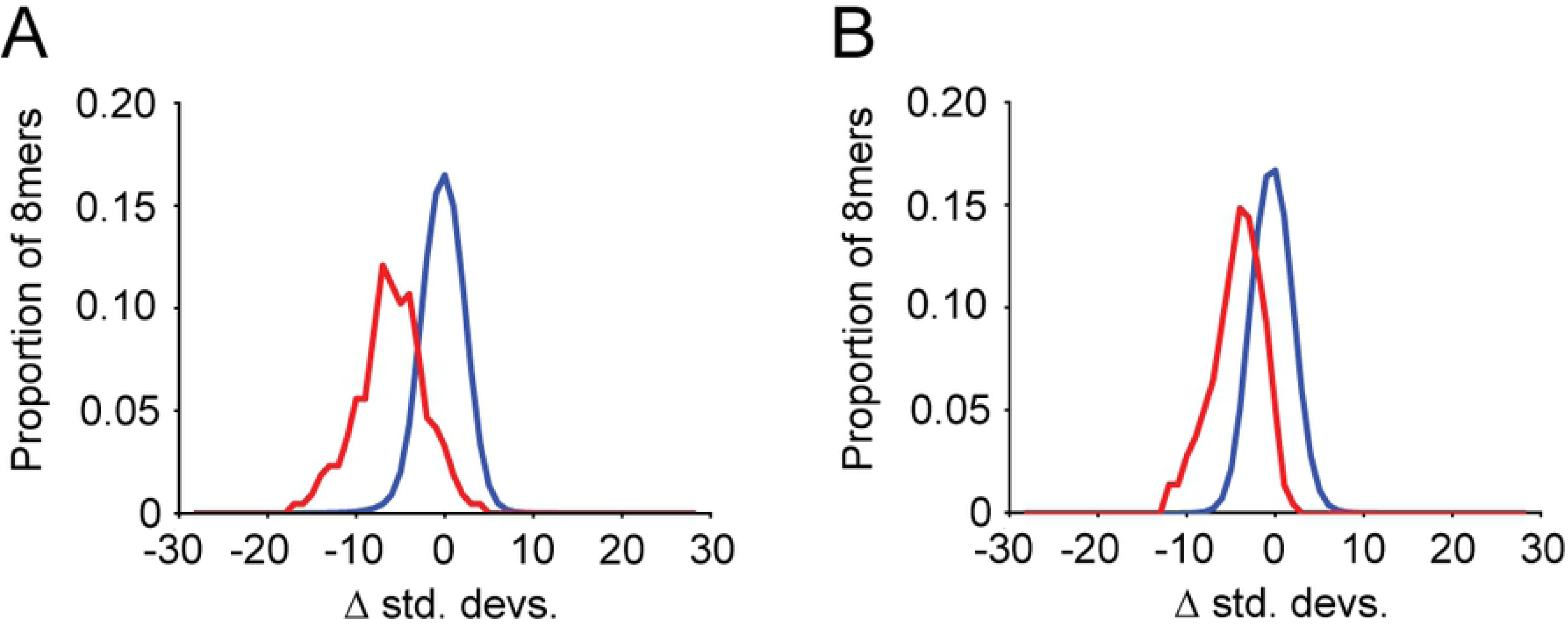
Analysis of microRNA target site loss and gain rates for deeply conserved miRNAs in drosophila species. Analysis as described in Fig 2.

**S3 Fig.**
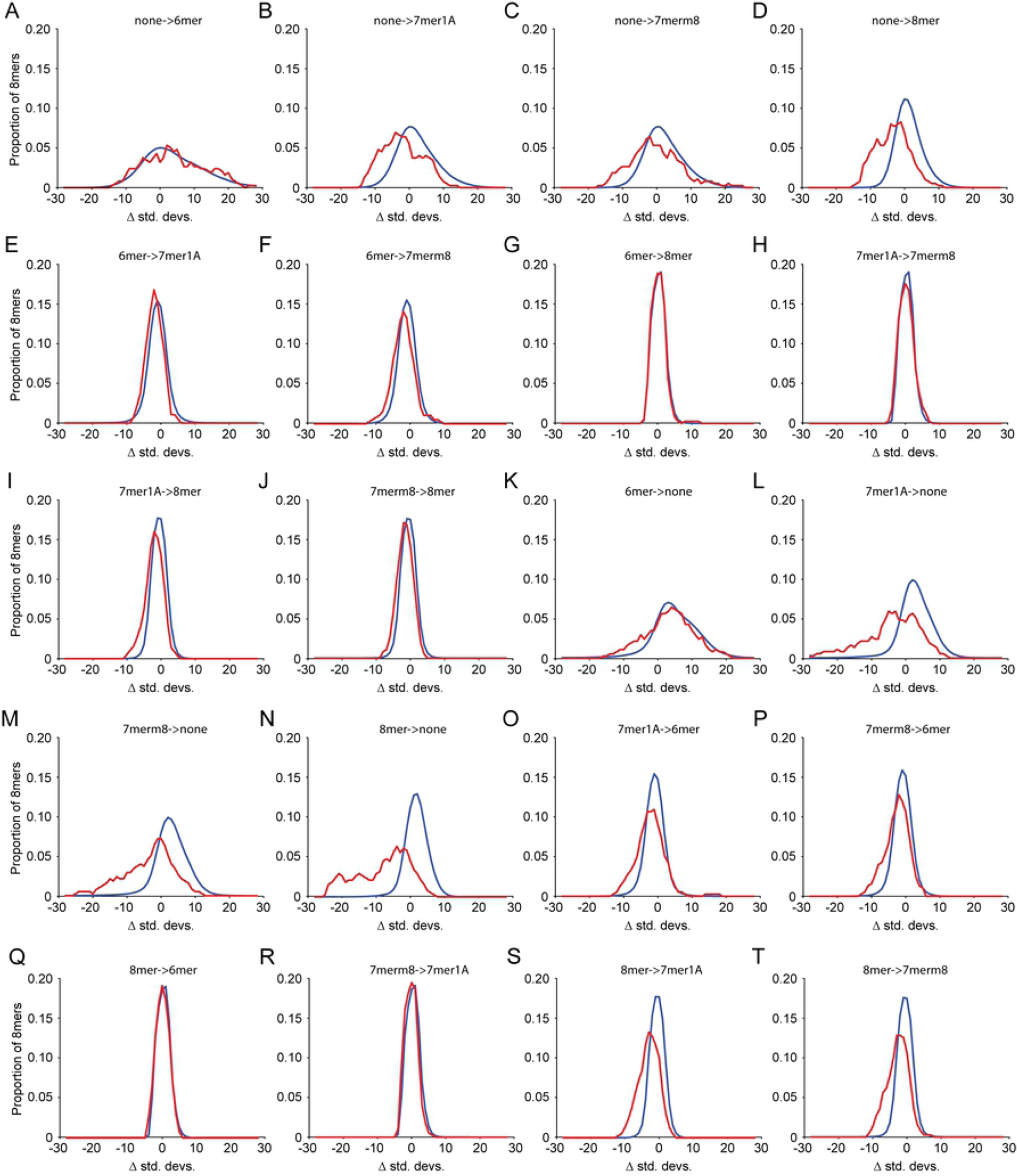
Interconversions that strengthen or weaken miRNA target sites are disfavored. Analysis of standard deviation metrics, calculated as in Fig 2, for target site interconversion events between the site types none, 6mer, 7merA1, 7merm8, and 8mer. Wilcoxon rank sum statistics for miRNAs (red line) relative to all 8mers (blue line) are captured as p-values and shown in Fig 6.

